# The JAK2-STAT pathway epigenetically regulates tolerized genes during the first encounter with bacterial antigens

**DOI:** 10.1101/2020.10.16.342717

**Authors:** Clara Lorente-Sorolla, Octavio Morante-Palacios, Antonio Garcia-Gomez, Laura Ciudad, Francesc Català-Moll, Adolfo Ruiz-Sanmartín, Mónica Martínez-Gallo, Ricard Ferrer-Roca, Juan Carlos Ruiz-Rodriguez, Damiana Álvarez-Errico, Esteban Ballestar

**Author notes:** Correspondence: E. Ballestar. These two authors contributed equally to the study (Clara Lorente-Sorolla, Octavio Morante-Palacios).

## Abstract

Microbial challenges, such as widespread bacterial infection, induce endotoxin tolerance. This state of hyporesponsiveness to subsequent infections is mainly displayed by monocytes and macrophages. Endotoxin tolerance is generally acquired following a septic episode. In this study, we investigated DNA methylation changes during the acquisition of *in vitro* tolerance. We identified a set of TET2-mediated demethylation events that are specific to toll-like receptor (TLR) 2 and TLR4 stimulation. Lipopolysaccharide (LPS)-specific demethylation occurs at genomic sites that have low accessibility in quiescent monocytes, concomitantly with the transcriptional activation of many inflammation-related genes, and they are enriched in binding motifs for several signal transducer and activator of transcription (STAT) family members. Indeed, STAT1, STAT3 and STAT5, elements of the JAK2 pathway, are phosphorylated in association with the acquisition of endotoxin tolerance. Inhibition of the JAK2 pathway impairs the activation of tolerized genes on the first encounter with LPS. This is evidence of a crucial role for this pathway in determining the initial response of these genes to bacterial antigens and provides a pharmacological target to prevent exacerbated responses, allowing regulated responses upon subsequent challenges. Finally, we assess the pathological relevance of the JAK2 pathway in monocytes from patients with sepsis.

## INTRODUCTION

Organisms are steadily exposed to threats from other species. Innate immune cells are the first line of host defense against invading pathogens. They activate the adaptive immune system to restore homeostasis and remove the infection (1). In innate immune cells, recognition of pathogen-associated molecular patterns (PAMPs) activates toll-like receptors (TLRs), triggering robust inflammatory responses (2). Different mammalian TLRs recognize distinct microbial ligands. For instance, Gram-negative bacterial lipopolysaccharide (LPS) specifically activates TLR4, while Pam3Cys-Ser-Lys4 (P3C), a synthetic analog of the triacylated N-terminal part of bacterial lipoproteins, binds TLR2.

In TLR2/4 signaling, inflammatory-related transcription factors such as nuclear factor kappa-light-chain-enhancer of activated B cells (NFκB), activator protein 1 (AP-1) and interferon regulatory factors (IRFs) are activated (3, 4). Moreover, as a consequence of the autocrine or paracrine activation of cytokine receptors such as interferon receptors, Janus kinases (JAKs) are, in turn, activated, which leads to the phosphorylation and recruitment of members of the signal transducer and activator of transcription (STAT) protein family. Phosphorylated STAT dimers are then translocated to the nucleus, where they bind to specific DNA sequences and initiate gene transcription (5). JAK/STAT activation is tightly regulated by members of the suppressors of cytokine signaling (SOCS) protein family, protein inhibitors of activated STATs (PIAS) and protein tyrosine phosphatases (PTPs) (6).

Monocytes and macrophages are highly plastic innate immune cells involved in phagocytosis and cytokine release. After stimulation, monocytes, being key mediators of the initial immune response, rapidly migrate to tissues (7). Monocytes and macrophages develop endotoxin tolerance, a state of hyporesponsiveness following a first challenge with microbial ligands (8). Tolerant cells show an altered immune response, characterized by lower production of pro-inflammatory cytokines (TNFα, IL-6, IL-1β, etc.) and higher levels of anti-inflammatory cytokines (IL-10, TGFβ, etc.) (9). Downregulation of major histocompatibility complex (MHC) class II molecules under endotoxin tolerance has also been observed (10). This immunosuppressive state can be achieved under pathological conditions, such as the state following a sepsis episode. Sepsis is characterized by a dysregulated inflammatory response driven by an infection. It is a potentially lethal disease, and, in particular, a major cause of death in intensive care units (11). Peripheral blood mononuclear cells (PBMCs) from patients with sepsis exposed *ex vivo* to LPS, express lower levels of TNFα, IL-6 and IL-1β compared with controls (12) and upregulated levels of IL-10 (13).

Endotoxin tolerance involves epigenetic mechanisms. For instance, LPS-treated murine monocytes display a group of tolerized genes with reduced gene expression and active histone marks after a second stimulus in comparison with untreated monocytes (14). Moreover, LPS-treated human monocytes show reduced levels of histone H3K27ac and H3K4me1 at promoters and enhancers of phagocytic and lipid metabolism genes(15). In addition, genome-wide analyses in human macrophages showed specific epigenetic signatures in H3K4me3, H3K27Ac and H3K4me1 histone marks for LPS tolerant cells (16). Studies in human monocytic THP-1 cells also revealed that H3K9 dimethylation of the promoters of the TNFα and IL-1β genes is responsible for their silencing during endotoxin tolerance (17). In fact, the histone H3K9 methyltransferase G9a, combined with HP1 and DNA methylation machinery, regulates TNFα gene expression (18).

Understanding the molecular and cellular mechanisms by which inflammatory response and endotoxin tolerance in LPS-treated monocytes are acquired and regulated could have clinical applications in new biomarkers or promising therapies. DNA methylation is potentially a significant mechanism in endotoxin tolerance, given its importance in monocyte/macrophage biology(19, 20) and its relative stability compared with other epigenetic marks. Recently, specific DNA methylation changes were found in monocytes from patients with sepsis associated with several clinical factors and functional features, supporting their relevance to the monocyte response to bacterial molecules and the course of the disease (21).

In this study, we integrate DNA methylation and transcriptome changes in an *in vitro* model of LPS-induced tolerance. We report a negative correlation between the two processes and a temporal sequence in relation to accessibility and active histone marks gains. Moreover, we show that JAK2-mediated pathways have a critical role in establishing the LPS-driven transcriptome and methylome remodeling, most probably due to engagement of the IFNγR upon autocrine/paracrine INFγ release, secondary to LPS activation, establishing a potential link with the regulation of genes that become tolerized. Our data on monocytes isolated from patients with sepsis indicate that JAK2/STAT pathway dysfunction may be relevant to endotoxin tolerance, and its pharmacological activation could improve the regulated expression of genes that become tolerized, providing a potential target to modulate their inflammatory response.

## MATERIAL AND METHODS

### Human samples

We selected and diagnosed patients with sepsis based on the criteria of the Third International Consensus Definitions for Sepsis and Septic Shock (Sepsis-3) (22). For each patient, we calculated the Sequential [Sepsis-related] Organ Failure Assessment (SOFA) score. The study included 4 patients with bacterial infections with SOFA scores ranging from 2 to 8. Patients were obtained from Vall d’Hebron University Hospital. Blood samples were collected within 12 h of sepsis diagnosis, which was confirmed using clinical and analytical data. The Committee for Human Subjects of Vall d’Hebron University Hospital (PR (ATR)122/2019) approved the study, which was conducted in accordance with the ethical guidelines of the 1975 Declaration of Helsinki. All samples were collected and processed in compliance with the guidelines approved by the local ethics committee. All participants (patients with sepsis and healthy controls) received oral and written information about the possibility that their blood would be used for research purposes before they gave their signed informed consent.

### CD14+ monocytes purification and culture

For *in vitro* experiments, we obtained buffy coats from anonymous donors via the Catalan Blood and Tissue Bank (CBTB). The CBTB follows the principles of the World Medical Association (WMA) Declaration of Helsinki. Before providing blood sample, all donors received detailed oral and written information and signed a consent form at the CBTB. PBMCs were isolated by density-gradient centrifugation using lymphocyte-isolation solution (Rafer, Zaragoza, Spain). Pure monocytes (MO) were then isolated from PBMCs by positive selection with magnetic CD14 MicroBeads (Miltenyi Biotec, Bergisch Gladbach, Germany). Purified monocytes were resuspended in Roswell Park Memorial Institute (RPMI) Medium 1640 + GlutaMAX™ (Gibco, Thermofisher) containing 10% human pooled serum (One Lambda, ThermoFisher Scientific Brand, West Hills CA, USA), 100 units/mL penicillin, and 100 mg/mL streptomycin. Monocytes were untreated (control), or treated with lipopolysaccharide (LPS) (10 ng/ml from *E. coli* O111:B4, Sigma-Aldrich, Darmstadt, Germany), and 10 μg/ml Pam3Cys (P3C) (InvivoGen San Diego, CA, USA). After 24 hours, monocytes were washed and left to rest for 3 days in medium supplemented with human pooled serum. Cells were then re-stimulated with LPS (10 ng/ml) and after 1 day, pelleted cells and supernatants were collected and stored until use.

For JAK2 inhibition of LPS monocytes (LPS+iJAK2 monocytes), cells were grown under the same conditions as mentioned above, and in the presence of fedratinib 500 nM (formerly known as TG101348, Santa Cruz Biotechnology), unless a different concentration is indicated.

### Cytokine measurements

The cytokines levels were measured from the cell culture supernatants using an enzyme-linked immunosorbent assay (ELISA), according to the manufacturer’s instructions (BioLegend, San Diego, CA, USA). In addition, the Pre-defined Human Inflammatory Panel LegendPlex™ (BioLegend) was used for the simultaneous analysis of 13 cytokines (CCL2, IFN-α2, IFN-γ, IL-1β, IL-6, IL-8, IL-10, IL-12p70, IL-17A, IL-18, IL-23, IL-33, and TNFα) related to inflammation in the same samples, in accordance with the manufacturer’s instructions.

### DNA methylation profiling and pyrosequencing

Infinium MethylationEPIC BeadChip (Illumina, Inc., San Diego, CA, USA) arrays were used to analyze DNA methylation. This platform allows over 850,000 methylation sites per sample to be interrogated at single-nucleotide resolution, and covers 99% of reference sequence (RefSeq) genes. The samples were bisulfite-converted using EZ DNA Methylation-Gold™ Kit (Zymo Research, CA, USA) and were hybridized in the array following the manufacturer’s instructions.

Each methylation data point was obtained from a combination of the Cy3 and Cy5 fluorescent intensities from the M (methylated) and U (unmethylated) alleles. For representation and further analysis, we used beta and M values. The beta value is the ratio of the methylated probe intensity to the overall intensity (the sum of the methylated and unmethylated probe intensities). The M value is calculated as the log2 ratio of the intensities of the methylated versus unmethylated probe. Beta values were used to derive heatmaps and to compare DNA methylation percentages from bisulfite-pyrosequencing experiments. For statistical purposes, the use of M values is more appropriate, because their degree of homoscedasticity fits better with linear model assumptions.

Bisulfite pyrosequencing was used to validate CpG methylation changes. DNA was isolated using a Maxwell^®^ RSC Cultured Cells DNA Kit (Promega). Bisulfite modification of genomic DNA isolated from monocytes was performed using an EZ DNA Methylation-Gold™ Kit (Zymo Research), following the manufacturer’s protocol. Primers for PCR amplification and sequencing were designed with PyroMark^®^ Assay Design 2.0 software (QIAGEN, Hilden, Germany). PCRs were performed with the IMMOLASE™ DNA polymerase PCR kit (Bioline), and the success of amplification was assessed by agarose gel electrophoresis. PCR products were pyrosequenced with the Pyromark Q24 system (QIAGEN).

### Quality control, data normalization, and detection of differentially methylated CpGs

Methylation array data were processed with the statistical language R using methods from the Bioconductor libraries minfi, lumi, and limma. Data quality was assessed using the standard pipeline from the minfi package(23). The data were Illumina-normalized, and beta and M values were then calculated. We excluded CpGs with overlapping SNPs. M values were used to build a linear model using the limma package in R, including the donor as a covariable.

In this study, we considered a probe to be differentially methylated if it had a methylation differential of 20% (Δβ ≥ 0.2) and when the statistical test was significant (false discovery rate (FDR) < 0.05).

### RNA purification and RNA-seq analysis

Total RNA of uncultured monocytes, LPS monocytes, LPS+iJAK2 monocytes, and untreated monocytes were isolated using a Maxwell^®^ RSC simplyRNA kit (Promega, Wisconsin, USA).

RNA-seq libraries were generated and paired-end sequenced in the Genomics Unit of the Centre for Genomic Regulation (CRG) (Barcelona, Spain).

Fastq files were aligned to the hg38 transcriptome using HISAT2 (24) with standard options. Reads mapped in proper pair, being primary alignments were selected with samtools (25). Reads were assigned to genes with FeatureCounts (26).

Differentially expressed genes were detected with DESeq2 (27). The donor was used as a covariable in the model. The Ashr shrinkage algorithm was applied and only protein-coding genes with abs(logFC) > 1 and FDR < 0.05 were selected as differentially expressed.

For representation purposes, Variance Stabilizing Transformation (VST) values and normalized counts provided by DESeq2 were used.

### Gene ontology over-representation analysis, Gene Set Enrichment Analysis and transcription factor enrichment analysis

Gene ontology (GO) over-representation of differentially methylated CpGs was analyzed using the Genomic Regions Enrichment of Annotations Tool (GREAT, version 4.0.4) (http://great.stanford.edu/public/html/), adopting the standard options (28) and using EPIC array CpGs as background. Enrichment is measured as the −log_10_ binomial FDR.

Gene Set Enrichment Analysis (GSEA) was performed from the LPS versus untreated, and LPS versus LPS-iJAK2 comparisons. Genes were ranked using this formula: −log_10_(FDR) * sign(log(FC)). As genesets collection, hallmarks (H) from the Molecular Signatures Database (MSigDB) were selected, adding the specified custom genesets. GSEA analysis and graphs were created with the ClusterProfiler (29) and enrichplot Bioconductor packages. Gene Ontology over-representation of upregulated and downregulated protein-coding genes was performed with ClusterProfiler, using all detected protein-coding genes as background.

We used the findMotifsGenome.pl command in the Hypergeometric Optimization of Motif EnRichment (HOMER) suite to look for motifs that are enriched in the target set relative to the background set (software v4.11) (30). It was used to identify enrichment of TF binding motifs in the 250bp-window upstream and downstream of the differentially methylated CpG sites. Annotated CpGs in the EPIC array were used as background.

### Accessibility and histone-mark profiling of differentially methylated CpGs

Using public data sets of ATAC-seq and H3K4me1 and H3K27ac ChIP-seqs of untreated and LPS-treated monocytes at 1, 4 and 24 hours (15), the accessibility and histone mark occupancy in the differentially methylated CpG genomic positions were calculated. Moreover, data of whole-genome bisulfite sequencing (WGBS) of the same reference were also utilized.

Graphs of the ATAC-seq and WGBS data were created with the deeptools toolkit (31).

For ChIP-seq data of H3K27ac and H3K4me1, bed files were downloaded from the BLUEPRINT portal (http://dcc.blueprint-epigenome.eu/). A file for each histone mark, cell type, and time point was used. Enrichment of these histone marks around the CpG positions (−3kb, 3kb) was studied applying a Fisher’s Exact Test to compare them with the background (EPIC array CpGs), dividing the studied region in tiles of 10 bp. The calculated odds ratio of each tile is represented.

### Methylation and expression association

Hypomethylated CpGs were associated with the nearest TSS using the annotatePeaks.pl command in the HOMER suite (30). After removing duplicate genes, expression of that gene set was studied using a public dataset of an RNA-seq time course (15) and our RNA-seq data.

### Quantitative reverse-transcription polymerase chain reaction (qRT-PCR)

Total RNA was isolated with a Maxwell^®^ RSC simplyRNA kit (Promega, Wisconsin, USA) and reverse-transcribed using a Transcriptor First Strand cDNA Synthesis Kit (Roche, Basel, Switzerland), in accordance with the manufacturer’s instructions. qRT-PCR was performed in triplicate using LightCycler^®^ 480 SYBR Green Mix (Roche). The standard double delta Ct method was used to determine the relative quantification of target genes, and values were normalized against the expression of endogenous control genes such as *RPL38*.

### Western blotting

Cytoplasmic and nuclear protein fractions were obtained using hypotonic lysis buffer (Buffer A; 10 mM Tris pH 7.9, 1.5 mM MgCl_2_, 10 mM KCl supplemented with protease inhibitor cocktail (Complete, Roche) and phosphatase inhibitor cocktail (PhosSTOP, Roche) to lyse the plasma membrane. Protein pellets were resuspended in Laemmli 1X loading buffer.

Proteins were separated by SDS-PAGE electrophoresis. Immunoblotting was performed on polyvinylidene difluoride (PVDF) membranes following standard procedures. Membranes were blocked with 5% bovine serum albumin (BSA) and blotted with primary antibodies. After overnight incubation, membranes were washed three times for 10 minutes with TBS-T (50 mM Tris, 150 mM NaCl, 0.1% Tween-20) and incubated for 1 hour with HPR-conjugated mouse or rabbit secondary antibody solutions (Thermo Fisher) diluted in 5% milk (diluted 1/10000). Finally, proteins were detected by chemiluminescence using WesternBright™ ECL (Advansta). The following antibodies were used:

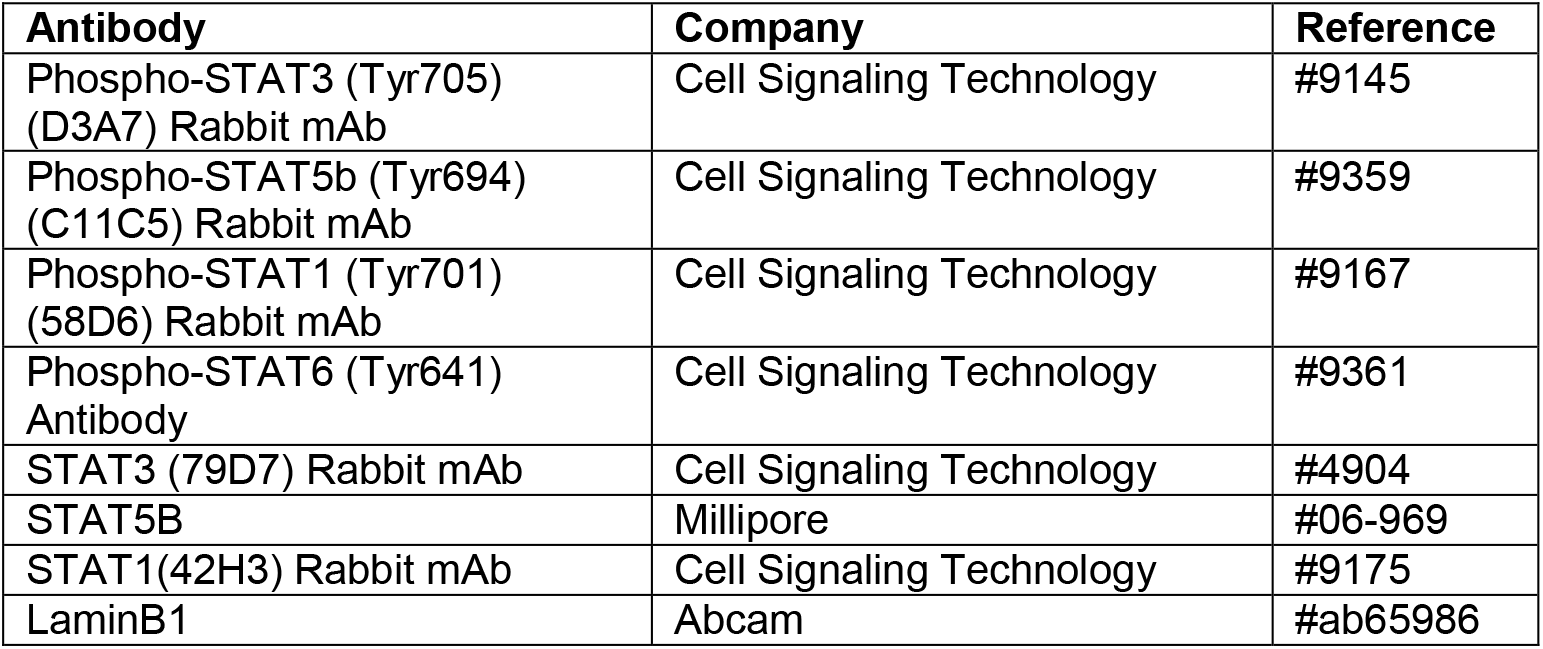

| Antibody | Company | Reference |
| --- | --- | --- |
| Phospho-STAT3 (Tyr705) (D3A7) Rabbit mAb | Cell Signaling Technology | #9145 |
| Phospho-STAT5b (Tyr694) (C11C5) Rabbit mAb | Cell Signaling Technology | #9359 |
| Phospho-STAT1 (Tyr701) (58D6) Rabbit mAb | Cell Signaling Technology | #9167 |
| Phospho-STAT6 (Tyr641) Antibody | Cell Signaling Technology | #9361 |
| STAT3 (79D7) Rabbit mAb | Cell Signaling Technology | #4904 |
| STAT5B | Millipore | #06-969 |
| STAT1(42H3) Rabbit mAb | Cell Signaling Technology | #9175 |
| LaminB1 | Abcam | #ab65986 |

### pSTAT1 flow cytometry

Peripheral blood mononuclear cells (PBMCs) were purified from blood samples of septic patients and healthy donors by density gradient centrifugation using lymphocytes isolation solution (Rafer, Zaragoza, Spain). PBMCs were counted and 5 million per condition were cultured in T25 flasks in Roswell Park Memorial Institute (RPMI) Medium 1640 + GlutaMAX™ (Gibco, Life Technologies, CA, USA) containing 2% Fetal Bovine Serum (FBS), 100 units/mL penicillin, and 100 mg/mL streptomycin, for 4 hours, with or without LPS (10 ng/ml from *E. coli* O111:B4, Sigma-Aldrich, Darmstadt, Germany). Cells were collected and stained with CD14, CD15, and pSTAT1 according to the BD Fixation/Permeabilization Solution Kit (#554714) and the antibodies manual.

| Antibody | Company | Reference |
| --- | --- | --- |
| Mouse Anti-pStat1 BV421 (pY701) | BD Biosciences | #562985 |
| Mouse IgG2a, $\kappa$ Isotype BV421 Control | BD Biosciences | #563464 |
| CD14 (APC) | Miltenyi Biotec | #130-091-243 |
| CD15 (FITC) | Miltenyi Biotec | #130-113-484 |

### Mann–Whitney U test and Student’s paired t test

Data were analyzed with Prism version 6.0 (GraphPad). Statistical analyses consisted of non-parametric Mann–Whitney U tests, to determine differences between pairs of separate groups, and Student’s paired t test, to compare the means of matched pairs of groups, except where indicated otherwise. The levels of significance were: *, *p* < 0.05; **, *p* < 0.01; ***, *p* < 0.001.

## RESULTS

### Stimulation of monocytes with LPS/P3C yields specific DNA demethylation

To investigate the potential role of DNA methylation during endotoxin tolerance, human CD14+ monocytes isolated from healthy donor blood were pre-incubated for 24 hours with lipopolysaccharide (LPS) or Pam3Cys (P3C), which signal through TLR4 and TLR2, respectively, and lead to a tolerized state. After the first stimulation, cells were washed and left to rest for 3 days in the presence of human pooled serum. Cells were then stimulated again with LPS for another 24 hours (Figure 1A). Monocytes maintained with only RPMI medium and serum for 1 + 3 days (Ø) were used as negative controls.

**Figure 1.**
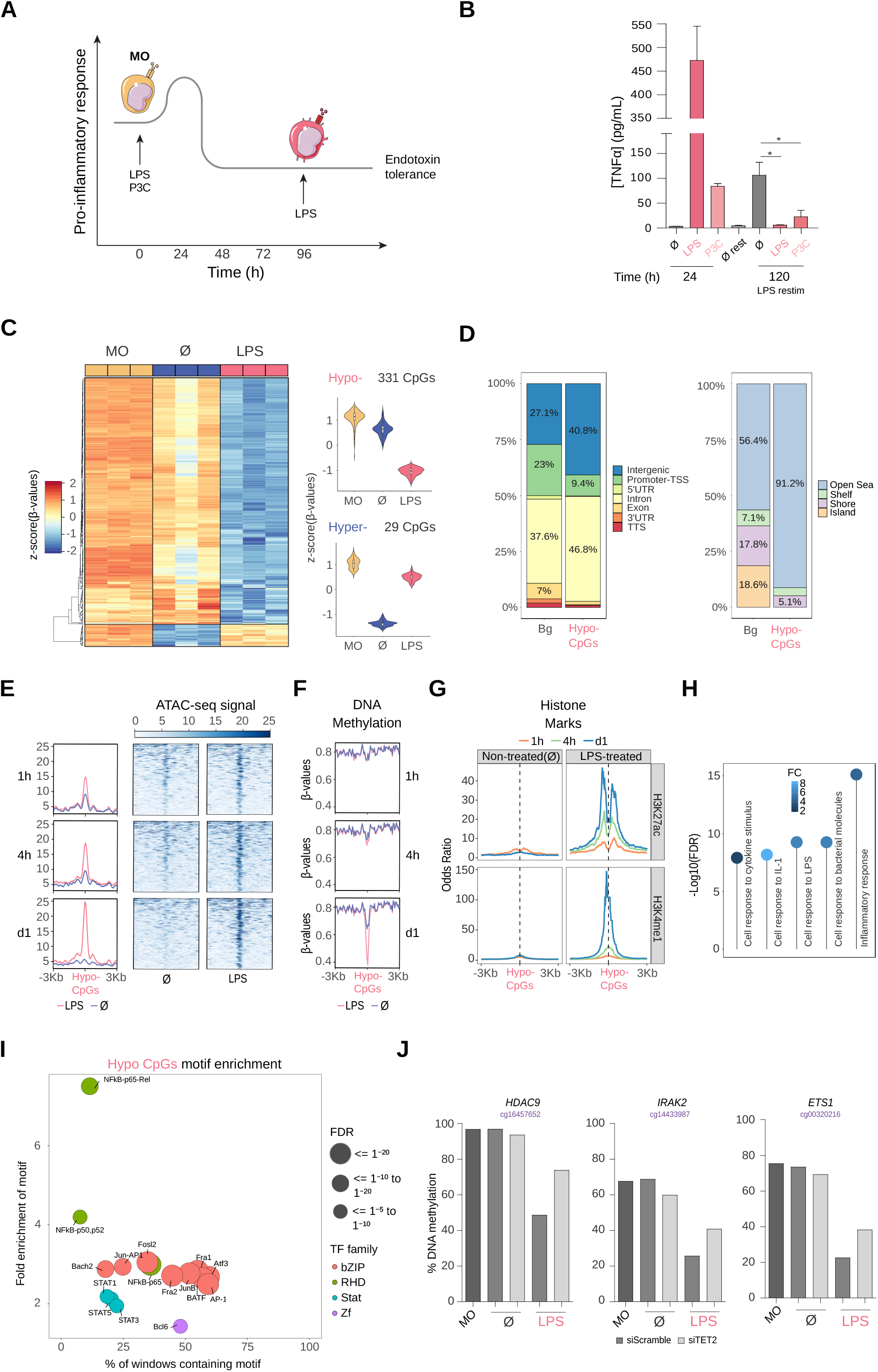
DNA methylation profile of LPS-treated human CD14+ monocytes. (A) Schematic diagram depicting *in vitro* experiments for endotoxin tolerance models. (B) Release of TNFα from CD14+ monocytes isolated from healthy donor blood samples treated with LPS (10 ng/ml, 24 hours) or P3C (10 μg/ml, 24 hours), then washed and rested (3 days) and treated again for 24 hours with LPS (10 ng/ml). These are all compared with untreated monocytes. Graphs show the mean ± SEM of three healthy donors. (C) DNA methylation heatmap of differentially methylated CpGs comparing untreated monocytes (Ø) with LPS monocytes (Δβ-≥ 0.2, adjusted *p* (FDR) < 0.05). Scaled β-values are shown, ranging from −2 (lower DNA methylation levels, blue) to +2 (higher methylation levels, red). On the right, violin plots of hypomethylated and hypermethylated clusters depicting normalized DNA methylation data. (D) Barplot presenting the percentages of different genomic features of hypomethylated CpGs in comparison with background (Bg) CpGs (left panel), and barplot of CpG island contexts (right panel). (E) Accessibility (ATAC-seq) data of hypomethylated CpGs after 1, 4 and 24 hours of monocyte culture with LPS (in red) or without treatment (Ø) (in blue). Public ATAC-seq data were used (15) (F) Methylation of hypomethylated CpGs after 1, 4 and 24 hours of monocyte culture with LPS (in red) or without treatment (Ø) (in blue). Public WGBS data were used (15). (G) ChIP-seq data of H3K27ac and H3K4me1 of LPS-treated and untreated monocytes were downloaded from the Blueprint database. Odds ratios were calculated for bins of 10 bp up to ±3000 bp around hypomethylated CpGs. CpGs annotated in the EPIC array were used as background. (H) GO (Gene Ontology) over-represented categories in the hypomethylated CpGs. The fold change relative to the background (EPIC array CpGs) and −log_10_(FDR) is shown. (I) Bubble scatterplot of TF binding motif enrichment for hypomethylated CpGs. The x-axis shows the percentage of windows containing the motif; the y-axis shows the magnitude of enrichment of the motif. Bubbles are colored according to TF family. FDR is indicated by bubble size (selected TF with FDR ≤ 1e^−05^). (J) DNA methylation measured by bisulfite pyrosequencing after TET2 silencing with siRNA (siTET2) in comparison with a control siRNA (siScramble).

Firstly, the levels of TNFα, which is a pro-inflammatory cytokine, were measured by ELISA to interrogate the acquisition of tolerance in monocytes treated under the previously described conditions. As expected, we observed that cells pre-exposed to LPS or P3C produced much lower levels of TNFα after the second LPS stimulation (Figure 1B).

We then obtained the DNA methylation profiles of the aforementioned samples, by hybridizing on Infinium MethylationEPIC bead arrays bisulfite-treated biological triplicates of monocytes, exposed for 24 hours to LPS or P3C, and after the 3-day resting period, with the matching untreated controls (4-days cultured monocytes; Ø) and uncultured monocytes at day 0 (MO).

Statistical analysis of the data revealed specific DNA methylation changes between LPS-/P3C-treated, untreated monocytes and uncultured monocytes (Supplementary Figure 1A). Methylation changes produced with LPS and P3C are comparable (Supplementary Figure 1B), suggesting that downstream events following engagement of TLR4 and TLR2 respectively (Supplementary Figure C) converge in the acquisition of similar DNA methylome changes.

Since LPS has a more intense effect than P3C (Supplementary Figure D), as reflected in the greater number of differentially methylated sites, but a very similar pattern (Supplementary Figure E), we focused on the analysis of changes between LPS-treated and untreated monocytes. We found 331 hypomethylated and 29 hypermethylated CpGs, with an FDR < 0.05 and a ΔB > 0.2. (Figure 1C). LPS-hypomethylated CpGs were located mostly in intergenic and intronic regions and were outside CpG islands (Figure 1D).

We then compared our DNA methylation data in relation to a time series (1, 4, 24 hours) of chromatin accessibility and DNA methylation datasets obtained under similar conditions (15). Interestingly, chromatin accessibility of the genomic positions bearing LPS-specific hypomethylated CpGs in LPS-treated monocytes started to increase very quickly (1 hour) and increased progressively over time (Figure 1E). In contrast, demethylation at such sites was only observed after 24 hours, indicating that accessibility precedes DNA methylation loss (Figure 1F). Similar to the changes in chromatin accessibility, histone marks characteristic of active enhancers (H3K4me1 and H3K27ac) also increased faster than DNA methylation changes (Figure 1G). In summary, the genomic composition and dynamics of LPS-demethylated sites indicate that a high proportion of methylation changes occur in regulatory regions, potentially controlling gene expression of phenotype-relevant genes. Moreover, these results suggest that LPS/P3C-driven demethylation requires a pioneer factor that can access closed chromatin following TLR4/TLR2 stimulation to enable, directly or indirectly, such specific demethylation.

Gene ontology (GO) over-representation analysis of hypomethylated CpGs in LPS-treated monocytes revealed the enrichment of functional categories associated with monocyte/macrophage cell biology and inflammation including cell response to cytokine stimulus, cell response to IL-1, cell response to bacterial molecules and inflammatory response (Figure 1H). In contrast, GO analysis of demethylated sites specific to untreated monocytes included regulatory categories such as regulation of IL-1 production, negative regulation of LPS signaling, and negative regulation of IL-12 production (Supplementary Figure 1F).

To detect potential transcription factors involved in the demethylation process, we performed a HOMER analysis. Hypomethylated CpGs of LPS-treated monocytes were enriched in transcription factor binding motifs relevant to inflammation such as NF-kB, the AP-1 family, and some members of the STAT family (STAT1, STAT5 and STAT3) (Figure 1I). In contrast, demethylated sites specific to untreated monocytes also contained motifs for NF-κB and AP-1, but not STAT family members (Supplementary Figure 1G), suggesting a specific role for this transcription factor family in the LPS-driven demethylation process most probably due to a second wave of chromatin changes triggered by IFN release a consequence of LPS-mediated monocyte response and the subsequent activation of the IFNR/JAK2/STAT axis that can in turn activate TET2 (32).

To validate the methylation array results, we selected some immune relevant genes, such as *CCL20*, *ETS1*, *HDAC9*, *IL24, IL2RA*, *IL36G* and *IRAK2*, from our methylation screening. Bisulfite pyrosequencing of these representative CpGs confirmed their specific demethylation in LPS and P3C samples (Supplementary Figure 1H).

We then downregulated ten-eleven translocation methylcytosine dioxygenase 2 (TET2) to determine whether this enzyme was involved in the demethylation processes observed under our conditions. TET2 has been implicated in catalyzing demethylation in other monocyte-related differentiation processes (19, 33). Using specific siRNAs, we achieved around 50% TET2 downregulation after 3 days of treatment (Supplementary Figure 1I). Under these conditions, demethylation of LPS-specific hypomethylated CpG sites was partially impaired (Figure 1J), demonstrating the involvement of TET2 in this process.

### LPS-driven gene expression changes are correlated with DNA methylation and are concomitant with STAT1, STAT3 and STAT5 activation

We performed RNA-seq with LPS-stimulated monocytes under the same conditions as before, including untreated and uncultured monocytes. LPS exposure induced upregulation of 1142 genes and downregulation of 1025 genes (logFC > 1, FDR < 0.05) in relation to untreated monocytes (Figure 2A). Apportioning the variance between the samples by principal component analysis (PCA), we observed that LPS-treated and untreated monocytes were separated along the axis of PC2, and these two along the PC1 axis with respect to uncultured monocytes (Supplementary Figure 2A).

**Figure 2.**
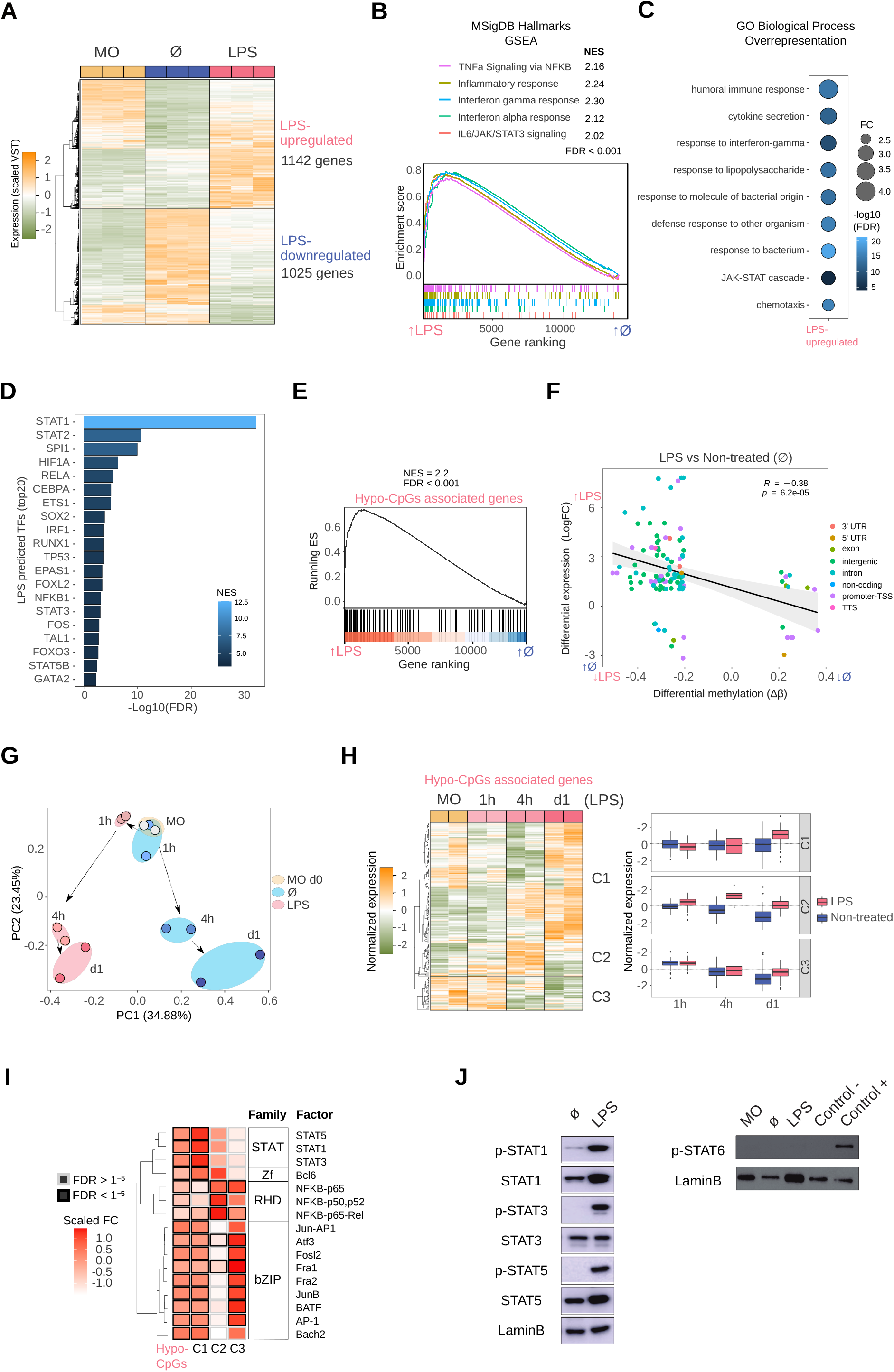
Gene expression, association with methylation and activation of JAK2-dependent STATs. (A) Gene expression heatmap of differentially expressed genes, comparing LPS monocytes with untreated (Ø) monocytes (LogFC > 1, FDR < 0.05). Scaled variance stabilizing transformation (VST) values are shown, ranging from −2 (lower gene expression level, green) to +2 (higher gene expression level, orange). (B) Gene set enrichment analysis (GSEA) of LPS versus untreated (Ø) monocytes, using MSigDB hallmarks (H) as gene sets. The running enrichment score and the normalized enrichment score (NES) are shown above the graph (FDR < 0.001). (C) Gene ontology (GO) over-representation of GO Biological Process categories. Fold change of LPS-upregulated genes relative to background and −log_10_(FDR) of the Fisher’s exact tests are shown. (D) Discriminant regulon expression analysis (DoRothEA) of LPS versus untreated (Ø) monocytes. NES and −log_10_(FDR) of transcription factors (TFs) enriched on the LPS side are shown (20 TF with the highest NES and FDR < 0.001). (E) Hypomethylated CpGs were associated with the nearest gene. The resulting gene set (Hypo-CpG-associated genes) was used in the GSEA of LPS versus untreated (Ø) monocytes. The running enrichment score and the normalized enrichment score (NES) are shown above the graph (FDR < 0.001). (F) DNA methylation of differentially methylated CpGs was correlated with gene expression of differentially expressed genes in the LPS versus untreated (Ø) monocyte comparison. Expression is represented on the y-axis as the LogFC, where higher values indicate higher levels of expression in LPS monocytes, and lower values indicate higher levels of expression in untreated monocytes. DNA methylation is depicted on the x-axis as Δβ, where lower numbers indicate lower levels of methylation in LPS monocytes, and higher numbers indicate lower levels of methylation in untreated monocytes. Points are colored according to their genomic context. A significant negative correlation between methylation and expression was observed (R = −0.38, p < 6.2e^−05^). (G) Hypo-CpG-associated gene expression was analyzed in time-course RNA-seq data from LPS-treated and untreated monocytes (15). Principal components 1 and 2 from a principal component analysis (PCA) of the expression data are shown. (H) Heatmap of Hypo-CpG-associated gene expression in a time-course RNA-seq, from LPS monocytes (left panel). The dendrogram can be considered to separate the sample into three clusters (C1, C2 and C3), depicting three distinct behaviors. Boxplot of normalized gene expression of genes in C1, C2 and C3 clusters, in LPS-treated (pink) and untreated (blue) monocytes. (I) Significant transcription factors of hypomethylated CpGs (Figure 1) were analyzed separately in the three clusters. Transcription factors are clustered by their position weight matrix differences. Scaled fold-change (FC) is represented as a color scale (a redder color indicates a greater change in relative to the background). The border of significant transcription factors (FDR < 1e^−5^) are colored black; non-significant borders are shown in grey. (J) Western blot of protein phosphorylated STAT1, total STAT1, phosphorylated STAT3, total STAT3, phosphorylated STAT5 and total STAT5 (left panel) and phosphorylated-STAT6 (right panel) in LPS-treated relative to untreated (Ø) monocytes. LaminB was used as loading control.

In the Gene Set Enrichment Analysis (GSEA), LPS-treated monocytes were enriched in significant inflammatory pathways such as TNFα signaling via NF-κB, inflammatory response, interferon gamma/alpha response, and IL6/JAK/STAT3 signaling (Figure 2B), indicating that the downstream targets of these signaling pathways had been transcriptionally activated. On the other hand, pathways depleted in LPS-treated monocytes included oxidative phosphorylation, MYC targets and fatty acid metabolism, as has also been described by others (Supplementary Figure 2B) (34, 35).

Similarly, GO over-represented categories in LPS-upregulated genes included terms such as cytokine secretion, response to interferon-gamma, response to LPS, and the JAK-STAT cascade (Figure 2C), with coincident categories with hypomethylated CpG GO categories (Figure 1H), whereas LPS-downregulated GO categories were related to metabolism (Supplementary Figure 2C).

To identify possible transcription factors leading to gene upregulation, we performed Discriminant Regulon Expression Analysis (DoRothEA), with which we identified several candidates coincident with those identified in our previous HOMER analysis of the LPS-specific DNA demethylation set: STAT (STAT1, STAT2, STAT3, STAT5B), NF-κB (RELA, NFKB1) and AP-1 (FOS) (Figure 2D). Additionally, in the set of downregulated genes, we found enrichment of transcription factors such as FOXP1, MYC and SREBF2 (Supplementary Figure 2D).

The similarities of methylation and expression GO categories and transcription factors potentially involved in the demethylation and upregulation reinforce the notion of their mutual relationship. In fact, LPS-upregulated genes are enriched in hypo-CpG-associated genes, as revealed by the GSEA analysis (Figure 2E). We also found a significant inverse correlation between DNA methylation and expression (Figure 2F).

Furthermore, using the expression data from the previously mentioned time-course study (1, 4 and 24 hours) in a similar model to ours (15), we monitored the expression of the LPS-hypomethylated CpGs-associated genes. Principal component analysis (PCA) revealed a divergent trajectory of LPS-treated and untreated monocytes (Figure 2G). The exposure of monocytes to LPS was sufficient to promote gene expression changes antagonistic to those in untreated monocytes, cultured without cytokines during the same period. Time-course expression analysis of LPS hypo-CpG-associated genes also revealed three main temporal clusters based on their dynamics (Figure 2H). The largest cluster (C1), showed an increase of gene expression at 24 hours concomitant with DNA demethylation (Figure 2H and Figure 1F). In contrast, cluster 2 (C2) presented an increase of gene expression prior to DNA demethylation, but coincident with the accessibility and enhancer-associated histone mark gains. Finally, cluster 3 (C3) showed a reduction in gene expression over time, in untreated and LPS monocytes, although the effect was less pronounced in the latter. These data imply that DNA methylation has different relationships with gene expression changes, possibly depending on the genomic context, and that most DNA methylation changes require previous chromatin remodeling, consistent with the timing observed for those changes to appear. Whereas gene upregulation is concomitant with DNA demethylation in C1, DNA demethylation occurs after gene upregulation in C2 and C3.

C1-, C2-, and C3-associated CpGs also had distinctive genomic features. C1-associated CpGs were predominantly located in introns and depleted in gene promoters (Supplementary Figure 2E), and were less accessible after 1 hour of LPS treatment. Their increasing accessibility was coincident with gene upregulation. By contrast, the maximum accessibility and gene expression of C2-associated CpGs both occurred by 4 hours of LPS treatment (Supplementary Figure 2F). Histone mark gains were also characteristic of each of these clusters, with only C1-associated CpGs showing a clear increase in both H3K4me1 and H3K27ac histone marks (Supplementary Figure 2G). The identification of such different states may reflect the dynamics of enhancer regulation during cell fate transitions following LPS stimulation, in which final cell identity may be stabilized in the end by the loss of methylation subsequent to chromatin remodeling, as it has been recently demonstrated in other myeloid cells (36).

HOMER analysis of C1-associated CpGs (with concomitant increase of expression and demethylation) revealed specific enrichment of STAT1, STAT3 and STAT5 binding motifs, which were not present in genes in C2 and C3 (Figure 2I), which points at the necessity of the activation of a second pathway such as IFNγR/JAK2/STAT in response to LPS triggered events in order to promote DNA methylation changes.

Since individual and combined DNA methylation and gene expression analysis suggested a specific role for the STAT transcription factor family, we studied the protein levels and phosphorylation of STAT1, STAT3 and STAT5 (JAK2 targets) in LPS-treated and untreated monocytes. We observed an increase in the protein level and phosphorylation of these three transcription factors (Figure 2J) in LPS-treated samples. In contrast, phosphorylated-STAT6 (JAK3 target), which did not appear in our HOMER analysis, was not detected in LPS monocytes, providing further evidence of the potential specific involvement of JAK2-related pathways.

### Inhibition of JAK2 partially prevents the activation of LPS-upregulated genes and tolerance-related genes

To investigate the potential role of JAK2 pathway in modulating tolerance-specific DNA methylation changes, we next performed previously described *in vitro* experiments in the presence of TG101348, a selective JAK2 inhibitor (iJAK2). CD14+ monocytes were treated with LPS (10 ng/ml) for 24 hours in the presence or absence of TG101348 (iJAK2) (500 nM). Monocytes were washed and left to rest for 3 days (with or without iJAK2) and after that, cells were stimulated again with LPS (24 hours). Control experiments without LPS were performed in parallel. We investigated phosphorylated protein levels using western blot and observed that JAK2-associated STAT phosphorylation was blocked by JAK2 inhibition, at 4 hours (left) and 96 hours (right) (Supplementary Figure 3A).

**Figure 3.**
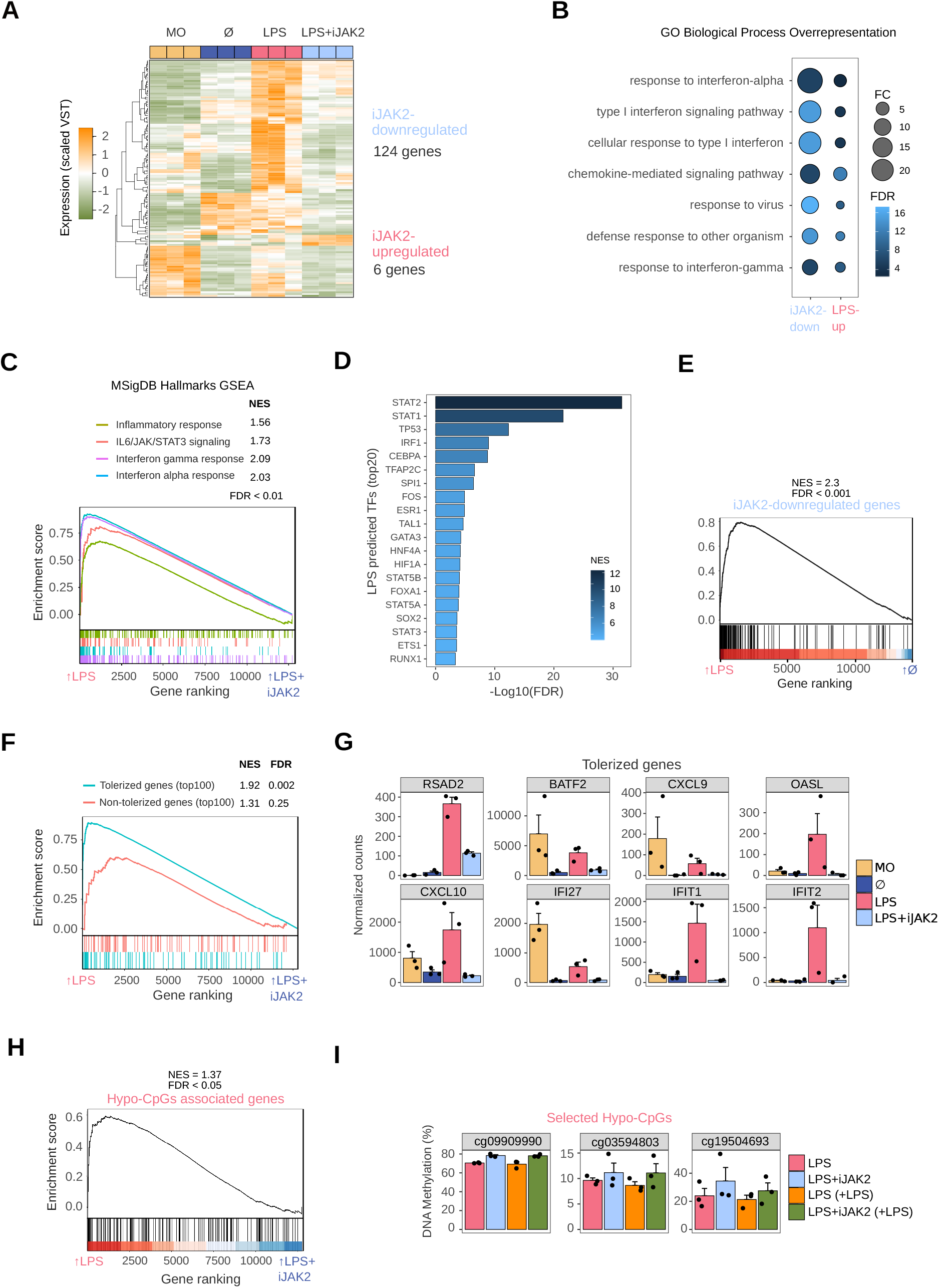
Role of JAK2/STAT in LPS monocytes gene expression and phenotype. (A) Gene expression heatmap of differentially expressed genes, comparing LPS with LPS+iJAK2 monocytes (LogFC > 1, FDR < 0.05). Scaled variance stabilizing transformation (VST) values are shown, ranging from −2 (lower gene expression level, green) to +2 (higher gene expression level, orange). (B) Gene ontology (GO) over-representation of GO Biological Process categories. Fold change of iJAK2-downregulated and LPS-upregulated genes relative to background and −log_10_(FDR) of the Fisher’s exact tests are shown. (C) Gene set enrichment analysis (GSEA) of LPS-treated versus LPS+iJAK2-treated monocytes, using MSigDB hallmarks as gene sets. The running enrichment score and the normalized enrichment score (NES) are shown above the graph (FDR < 0.001). (D) Discriminant regulon expression analysis (DoRothEA) of LPS-treated versus LPS+iJAK2-treated monocytes. NES and −log_10_(FDR) of transcription factors (TF) enriched on the LPS side are shown (20 TF with the highest NES and FDR < 0.001). (E) GSEA of LPS-treated versus untreated (Ø) monocytes using the iJAK2-downregulated genes as the gene set. The running enrichment score and the normalized enrichment score (NES) are shown above the graph (FDR < 0.001). (F) Gene Set Enrichment Analysis (GSEA) of LPS vs LPS+iJK2 monocytes using the top100 tolerized genes (blue) or the top100 non-tolerized genes (pink) as gene sets. Running enrichment score is represented and the normalized enrichment score (NES) and FDR are shown above. (G) Selected examples of iJAK2-downregulated genes. Bar plots show the mean ± SEM (standard error of the mean) of normalized counts. (H) GSEA of LPS-treated versus LPS+iJAK2-treated monocytes using the Hypo-CpG-associated genes as the gene set. The running enrichment score and the normalized enrichment score (NES) are shown above the graph (FDR < 0.05). (I) DNA methylation percentage obtained from pyrosequencing of three selected CpGs from the hypo-CpG group in LPS monocytes, before and after a second LPS stimulus, and with or without iJAK2 treatment.

To examine the effects of JAK2 inhibition on innate immune response, we measured the production of TNFα and IL-10 (Supplementary Figure 3B). JAK2 inhibition resulted in decreased production of TNFα and increased IL-10 production after a second LPS stimulus (Supplementary Figure 3B). This further reduction in TNFα production is consistent with the role of JAK2 in LPS-mediated inflammatory response (37).

To test the effects of JAK2 inhibition on the transcriptome, we performed RNA-seq of biological triplicates under the previously described cell conditions (LPS-treated monocytes, LPS+iJAK2-treated monocytes, untreated monocytes, and uncultured monocytes). JAK2 inhibition resulted in downregulation of 124 genes (iJAK2-downregulated genes), whereas only 6 genes were upregulated (LogFC > 1, FDR < 0.05) (iJAK2-upregulated genes) (Figure 3A).

The over-represented GO categories among the genes downregulated upon JAK2 inhibition (Figure 3B) were very similar to those found in LPS-upregulated genes (Figure 2C), including categories related to response to interferon-alpha, interferon-gamma, defense response to other organisms, etc. However, we found a generally higher magnitude change and a lower FDR in the iJAK2-downregulated genes than in the LPS-upregulated genes, suggesting specific and stronger enrichment in interferon-related genes.

In this respect, the GSEA analysis revealed four hallmarks downregulated in LPS+iJAK2 cells: inflammatory response, IL6/JAK/STAT3 signaling, interferon gamma response and interferon alpha response (Figure 3C). These categories were previously associated with LPS-treated monocytes (Figure 2B).

DoRothEA analysis showed several putatively related transcription factors depleted in LPS+iJAK2 monocytes, including STAT1, STAT2, STAT3, STAT5A and STAT5B (Figure 3D). NF-κB was not identified by either the DoRothEA or the GSEA analysis, in contrast to the LPS-associated hallmarks and transcription factors (Figure 2B, Figure 2D).

A relationship, as suggested by previous data, was found between iJAK2-downregulated genes and LPS-upregulated genes. In a GSEA, iJAK2-downregulated genes were strongly enriched in LPS-upregulated genes side in comparison with Untreated(Ø) monocytes (Figure 3E).

We also found a link between iJAK2-downregulated genes and tolerized genes. Using a public dataset (15) of gene expression (log2RPKM) of LPS-treated and untreated monocytes before and after a second LPS stimulus, we defined a ‘score’ of tolerization as *(Untreatedreexposure – Untreated) – (LPSreexposure – LPS)*, where a positive score means that the gene is tolerized and a negative score that is non-tolerized. We then compiled a top100-tolerized gene-set comprising the 100 genes with the highest scores and a top100-non-tolerized gene-set comprising the 100 genes with the lowest scores. We performed a GSEA analysis of both gene-sets with the comparison LPS/LPS+iJAK2 and found specific significant enrichment of the tolerized genes but not of the non-tolerized genes among the iJAK2-downregulated genes (Figure 3F). Some examples of tolerized-gene expression are shown in Figure 3G, where the direct inhibition of their transcriptional activation by JAK2 inhibition is depicted. Notably, these genes are also involved in the IFNα or IFNγ response.

Given the correlation between our DNA methylation and expression datasets (Figure 2F), the enrichment of STAT1/3/5 in the individual and combined data, and the effects of JAK2/STAT inhibition in the regulation of some LPS-upregulated genes (Figure 3F), we studied the effects of the inhibition of the JAK2/STAT pathway on hypomethylated CpG-associated genes. The GSEA of LPS versus LPS+iJAK2 monocytes revealed a small but significant enrichment in the LPS side, suggesting that genes related to CpGs hypomethylated with LPS are, at least partially, under the transcriptional control of the JAK2/STAT pathway (Figure 3H).

To determine whether the JAK2/STAT pathway is related to the LPS-specific DNA demethylation process we carried out pyrosequencing of three selected CpGs, with and without a second LPS stimulus (Figure 3I). A general trend of demethylation blockage was found, although only one CpG (cg09909990) had a value of p < 0.05. The fact that iJAK2 did not fully inhibit the phosphorylation of STATs, and the possible redundancy of other JAK/STAT pathways could explain the heterogeneity in the triplicate.

### STAT1 phosphorylation is reduced in monocytes from septic patients after an LPS stimulus

Given the relevance of JAK2/STAT pathways secondary to LPS response in monocytes, and to the transcriptional regulation of tolerized genes, we obtained monocytes from patients with sepsis to test whether the STAT activation was altered.

We incubated PBMCs from patients with gram negative bacterial sepsis and healthy donors in RPMI (2% FBS) for 4 hours, in the presence of LPS (10 ng/mL) and in its absence. Since activated neutrophils can contaminate the PBMC section of Ficoll and can express CD14, we adopted a gating strategy to analyze only the intracellular pSTAT1 signal of CD14+ CD15- cells (Q1) (Figure 4A).

**Figure 4.**
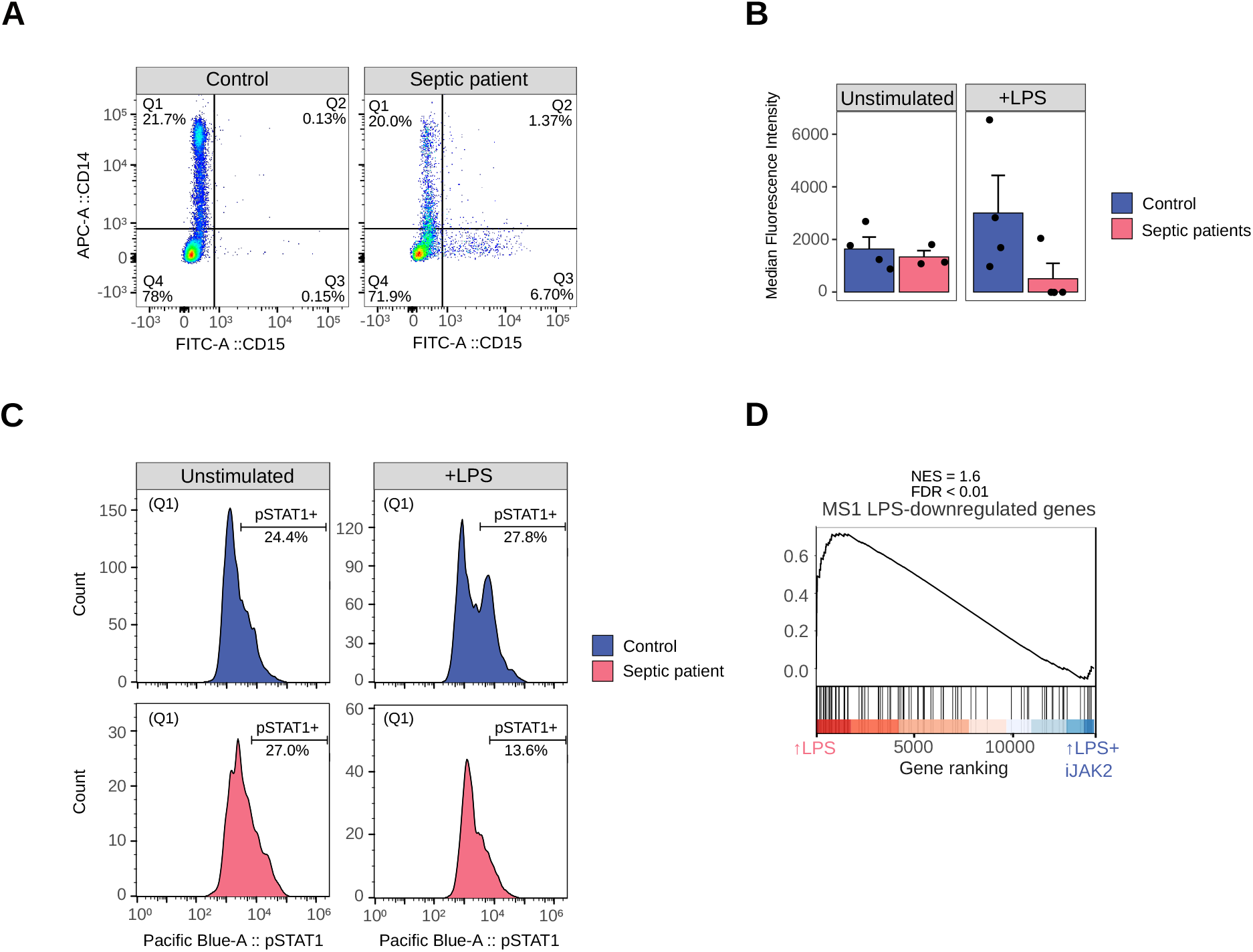
STAT1 phosphorylation in monocytes from healthy donors and septic patients with or without an LPS stimulus. (A) Example of the strategy adopted to gate only monocytes. CD14+CD15-(Q1) are mostly monocytes, whereas some CD15+ cells (neutrophils) can also express CD14 (Q2). (B) Median fluorescence intensity (pSTAT1) of monocytes (Q1) from septic patients (pink) and healthy donors (blue), with and without an LPS stimulus. (C) Example of a pSTAT1 signal histogram from a healthy donor (blue) and a septic patient (pink), with and without an LPS stimulus. (D) GSEA of LPS-treated versus LPS+iJAK2-treated monocytes using MS1 LPS-downregulated genes (38) as gene set. The running enrichment score and the normalized enrichment score (NES) are shown above the graph.

No differences were found between healthy donors and septic patients in the basal (unstimulated) signal of pSTAT1. However, after the LPS stimulus, pSTAT1 in healthy donors was increased, as expected, but decreased in septic patients (Figure 4B,4C).

To provide more insight into the JAK2-STAT1 pathway relevance in sepsis, we used expression data from a single-cell study of PBMCs from septic patients (38). They defined a specific monocyte subset associated with sepsis (MS1), showing a tolerized phenotype. After isolating that population and stimulating with LPS in vitro, some genes were downregulated (MS1 LPS-downregulated genes). We performed a GSEA comparing our LPS versus LPS+iJAK2 monocytes expression datasets revealing enrichment of the MS1 LPS-downregulated gene-set in the LPS side (Figure 4D). This result is consistent with the decrease of STAT1 activation in monocytes from septic patients after an LPS stimulus (Figure 4B) and shows that the inhibition of JAK2 in monocytes can recapitulate partially the expression changes produced in septic monocytes after a second immune challenge.

This result suggests that the JAK2-STAT1 pathway which is altered in septic patients, and partially their tolerized phenotype, causes reduced levels of pSTAT1 after a second immune challenge with LPS, potentially leading to abnormal methylation patterns that could partially drive the dysfunctional monocyte response in septic monocytes. Our results may explain the previously reported partial restoration of leukocytic function observed in septic patients after therapy with recombinant interferon-γ (39).

## DISCUSSION

Our results indicate that TLR4/TLR2 stimulation induces specific TET2-dependent demethylation in monocytes, accompanying the acquisition of endotoxin tolerance. A majority of LPS (TLR4)-specific changes in DNA methylation are concomitant with upregulation of inflammatory-related genes, and these involve the JAK2/STAT pathway. Inhibition of the JAK2 pathway in this *in vitro* model of endotoxin tolerance impairs the upregulation of genes that become tolerized following a first encounter with bacterial LPS.

Our analysis revealed several CpG sites that become demethylated after TLR4/TLR2 stimulation, many of which are associated with inflammatory genes. Examples include the chemokine *CCL20*, which has antimicrobial activity (40), *IL36G*, member of the IL-1 cytokine family, and the inflammatory cytokine *IL-24* (41). Other demethylated CpGs are associated with genes encoding the transcription factor *ETS1* and the molecule *IRAK2*, an essential regulator for IL-1R and TLR signaling (42). The majority of CpGs undergoing LPS-driven demethylation, are preceded by increases in chromatin accessibility and H3K4me1/H3K27ac gains and correlate with transcriptional activation of the associated genes. This temporal uncoupling between chromatin accessibility, transcription and DNA methylation in terminally differentiated myeloid cells, has also been recently demonstrated by simultaneously assessing chromatin conformational changes and DNA methylation in a genome-wide manner on the same population of DNA molecules (ATAC-Me technique), supporting the idea of DNA methylation as a required event in during enhancer activation that underlies cell state transitions (36). Moreover, transcription factors can directly recruit DNMTs or TET enzymes and influence gene expression (43). Our analysis revealed that binding motifs of STAT transcription factors family specifically associate with the observed LPS-specific hypomethylated CpGs and expression changes. In fact, the identified binding motifs correspond to STAT1, STAT3 and STAT5, which are phosphorylated by JAK2.

JAK/STAT signaling is not directly downstream of TLR4/TLR2 receptors, and its activation requires the production of other molecules, such as IFNγ or IL-6, in order to activate the pathways autocrinally or paracrinally through their receptors (6, 44). This could explain why STAT factors are specifically associated with C1-associated CpGs, whose associated genes are ‘late responsive’ in comparison with others, upregulated prior to demethylation (C2- and C3-associated CpGs). These CpGs are enriched in NF-κB and AP-1, directly downstream of TLR4/TLR2 signaling.

In our analysis, JAK2/STAT inhibition appeared to accentuate the tolerant phenotype of monocytes. JAK2 inhibition also reduces the expression of tolerized genes (15) following the first encounter with LPS, suggesting that these genes are under the transcriptional control of JAK2/STAT. In fact, several of these genes are also involved in the IFNα or IFNγ response.

In contrast, JAK2 inhibition only partially interferes with LPS-specific DNA demethylation, supporting the existence of a complex genomic regulatory network in which JAK2/STAT plays a role in LPS-specific demethylation, but also involves the participation of additional pathways.

We can speculate whether the observed effects of this JAK2 inhibition could be countered by direct activation with, for instance, IFNγ, which could reduce the acquired endotoxin tolerance and restore the expression of some tolerized genes. In this respect, demethylation has been previously associated with the IFNγ pathway, through STAT1 stimulation, binding to TET2 and recruitment to specific sites in the genome (32). This suggests that the direct modulation of this pathway may have more widespread effects on DNA methylation and expression. In fact, the metabolic effects of endotoxin tolerance are known to have been reverted in monocytes through IFNγ treatment, whose signaling pathway is JAK2-dependent (39).

The link between tolerized genes and downregulated genes upon JAK2 inhibition indicates that JAK2-dependent signaling pathway dysfunction may contribute to the acquisition of endotoxin tolerance.

Since monocytes from septic patients have been exposed to bacteria before they are extracted, they have a reduced immune response in a second immune challenge with LPS due to the endotoxin tolerance. We have shown that monocytes from septic patients exhibit a lower level of STAT1 phosphorylation after an immune challenge with LPS, compared with monocytes from healthy donors. This lower level of pSTAT1 due to a reduced activity of JAK2, could explain such a phenotype. In fact, IFNγ, whose receptor signals through JAK2, is believed to partially rescue the endotoxin tolerance phenotype in human monocytes (39), and IFNγ is sometimes used in sepsis treatment to improve immune host defense (45), perhaps activating JAK2 more intensively.

Taken together, our results demonstrate an important role of the JAK2/STAT pathway in the monocyte LPS response, which contributes not only to the DNA methylation, but also to gene expression remodeling. We also show that dysfunction in this pathway can be related to the phenomenon of endotoxin tolerance, in an *in vitro* model and in *ex vivo* monocytes from septic patients.

## Supplementary Figure Legends

**Supplementary Figure 1. DNA methylation profile of P3C-treated human CD14+ monocytes.** (A) DNA methylation heatmap of differentially methylated CpGs obtained with all the possible contrasts between the represented groups (Δβ-≥ 0.2, adjusted *p* (FDR) < 0.05). Scaled β-values are shown, ranging from −2 (lower DNA methylation levels, blue) to +2 (higher methylation levels, red). (B) Principal component analysis (PCA) of the CpGs shown in Supplementary Figure 1A. Principal components 1 and 2 are shown. (C) Simplified scheme of TLR4 and TLR2 signaling pathways. (D) Venn diagram of differentially methylated CpGs of LPS-treated versus untreated monocytes (LPS) and P3C-treated versus untreated monocytes (P3C). (E) Violin plots of LPS hypomethylated, LPS hypermethylated and P3C hypomethylated CpGs depicting normalized DNA methylation data. (F) GO (gene ontology) over-represented categories in M2 cluster CpGs. Fold change relative to background (EPIC array CpGs) and −log_10_(FDR) are shown. (G) Bubble scatterplot of TF binding motif enrichment for M2 cluster CpGs. The x-axis shows the percentage of windows containing the motif and the y-axis shows the magnitude of enrichment of the motif. Bubbles are colored according to the TF family. The FDR is indicated by the bubble size (selected TF with FDR ≤ 1e^−05^). (H) Percentage methylation level of some selected LPS-hypomethylated CpGs as obtained by bisulfite pyrosequencing in the LPS and P3C samples. (I) qRT-PCR analysis to validate the downregulation of TET2 by siRNA (siScramble used as control). Data are normalized relative to the RPL38 gene.

**Supplementary Figure 2.** (A) Principal component analysis (PCA) of the 500 most variable genes. Principal components 1 and 2 are shown. (B) Gene set enrichment analysis (GSEA) of LPS-treated versus untreated (Ø) monocytes, using MSigDB hallmarks (H) as gene sets. Gene sets enriched in untreated (Ø) monocytes are shown. The running enrichment score and the normalized enrichment score (NES) are shown above the graph (FDR < 0.001). (C) Gene ontology (GO) over-representation of GO biological process categories. Fold change of LPS-downregulated genes relative to background and −log_10_(FDR) of the Fisher’s exact tests are shown. (D) Discriminant regulon expression analysis (DoRothEA) of LPS-treated versus untreated (Ø) monocytes. NES and −log_10_(FDR) of transcription factors (TF) enriched on the Ø side are shown (FDR < 0.05). (E) Barplot of genomic features as percentages of C1-, C2- and C3-associated CpGs (Figure 2H) compared with total hypo-methylated CpGs. (F) Accessibility (ATAC-seq) data of CpGs associated with C1 (in red), C2 (in blue) and C3 (in green) after 1, 4 and 24 hours of monocyte culture with LPS. Public ATAC-seq data were used (15) (G) ChIP-seq data of H3K27ac and H3K4me1 of LPS-treated monocytes were downloaded from the Blueprint database. Odds ratios were calculated with C1-, C2- and C3-associated CpGs, using EPIC array CpGs as background. Odds ratio and −log_10_(FDR) obtained from Fisher’s exact tests are shown.

**Supplementary Figure 3.** (A) Western blot of JAK2-dependent STAT proteins in LPS-treated and untreated (Ø) monocytes, with and without JAK2 inhibitor (iJAK2) after 4 hours (left) or 96 hours (right) of treatment. LaminB was used as the loading control. (B) TNFα and IL-10 levels measured in cell supernatants to assess the tolerance state after JAK2 inhibition.

