## Supplementary figures and images for "The JAK2-STAT pathway epigenetically regulates tolerized genes during the first encounter with bacterial antigens"

### Supp. Figure 1

#### Supplementary Figure 1

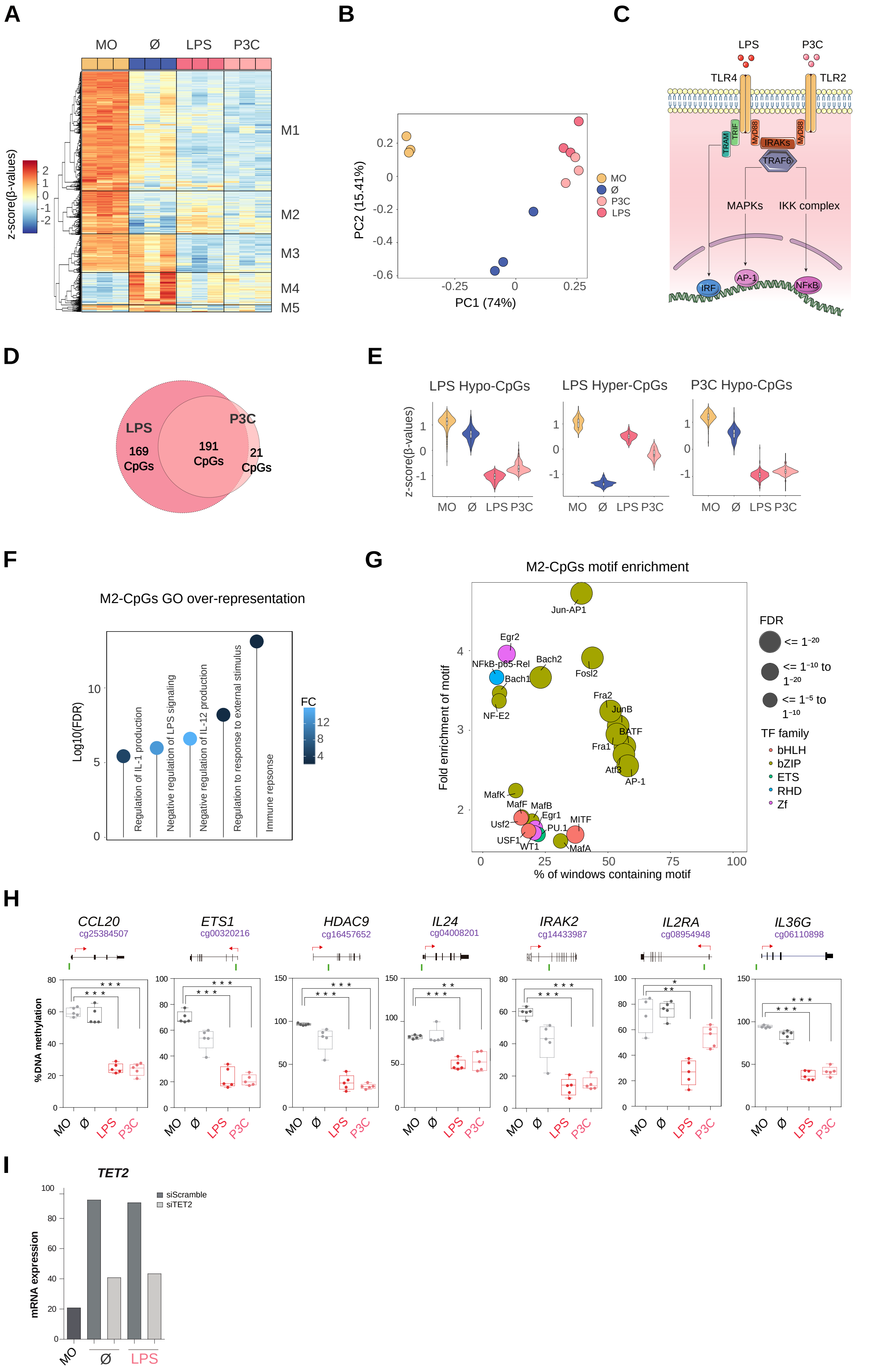

### Supp. Figure 2

#### Supplementary Figure 2

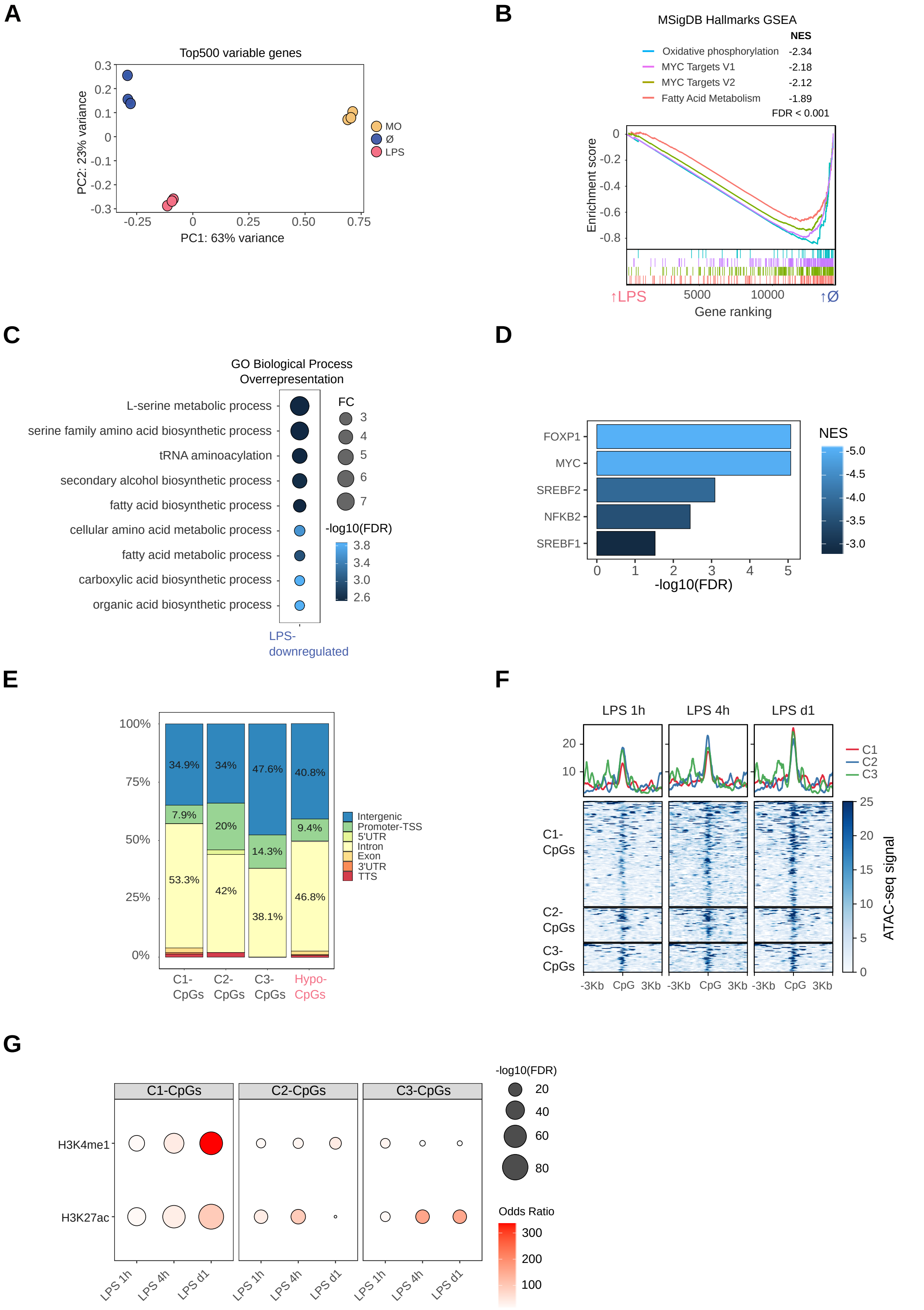

### Supp. Figure 3

#### Supplementary Figure 3

A

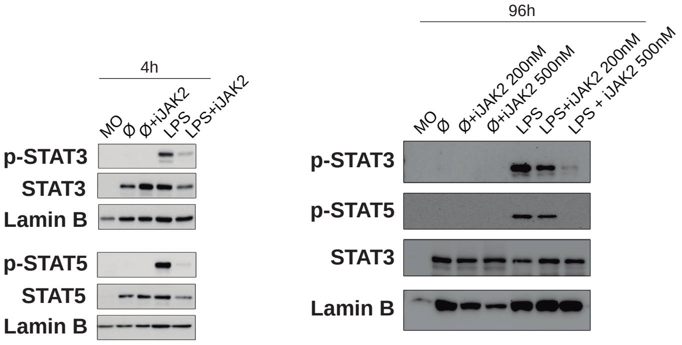


B

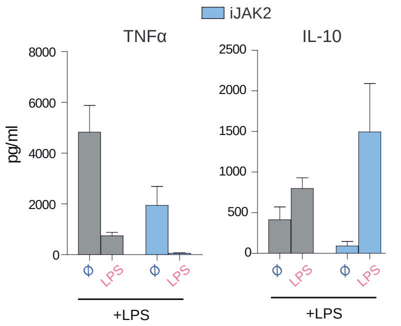
